# A meta-analysis of operational interactions between pinnipeds and fisheries

**DOI:** 10.1101/2022.11.24.517843

**Authors:** J Jackson, W Arlidge, R Oyanedel, KJ Davis

## Abstract

The global recovery of pinniped populations is a conservation success. However, pinniped population recovery has increased human-wildlife conflict with fisheries, an issue often reported and requiring management, but one that lacks global synthesis. We conduct a meta-analysis to estimate the impacts of operational interactions (specifically, lost catch) between pinnipeds and fisheries. Where quantifiable interactions are reported (n = 36), on average fishers have a 33.7% chance of interacting with pinnipeds on any given fishing day, and 13.8% lost catch. We find a large degree of heterogeneity between studies, with some fisheries experiencing much more negative interactions than others. Specifically, smaller-scale fisheries using nets are up to twice as likely to have negative interactions and lose up to five times more catch compared to large-scale fisheries. We conclude that pinniped-fishery conflict is a substantial global issue, but its impacts are not uniform. To successfully manage long-term coexistence between pinnipeds and humans, explicit data quantifying operational interactions is required. Population recoveries can have unintended consequences for fisheries, and management of ecological, social and economic outcomes is needed for long-term coexistence.

**Teaser:** Pinniped population recoveries have led to significant impacts on fisheries, but small-scale fisheries lose out most.

## Introduction

Conflict between marine predators and fishers is a long-standing global issue that has widespread impacts on both conservation outcomes and livelihoods (*1*). Fisheries are a key global industry, with a spatial extent over four times greater than terrestrial agriculture and a sector first sale value of USD 406 billion in 2019 (*2, 3*). However, fishing activity inevitably attracts marine predators, particularly cetaceans and pinnipeds, which have prey species in common with fishers (*4*). Pinnipeds, a suborder which includes the seals, sea lions and walruses, are prominently reported in interactions with fisheries, largely due to marine protection and population recoveries (*5, 6*). Historically, fishers managed interactions with pinnipeds through systematic and commercial hunting, resulting in average population declines of pinnipeds of over 70% relative to historic baselines until the mid-20^th^ century (*7*). In the 20^th^ century the systematic eradication of pinnipeds coupled with widespread overfishing resulted in many species being threatened with extinction or going extinct (*8*–*10*). Following successful conservation management initiatives in the 21^st^ century, such as the landmark US Marine Mammal Protection Act of 1972, many pinniped species are now recovering (*5, 11*). Magera et al. (*6*) estimated that 44-58% of pinniped species have seen significant increasing population trends since the end of the 20^th^ century. One compelling example is the northern elephant seal (*Mirounga angustirostris*), which had >48,000 births in the United States of America in 2010 compared to 132 in 1958 (*12*).

However, pinniped population recovery presents novel conservation challenges as managers are faced with the triple bottom line, striving to balance ecological, social, and economic objectives (*13, 14*). One unintended consequence of pinniped recovery worldwide is a resurgence in the reporting of fishery-pinniped conflict, most commonly through pinniped depredation of fishery catch or damage to fishing gear (i.e., operational interactions) (*4, 15, 16*). To achieve ecosystem-based marine management, a key goal of marine organisations worldwide (*17*), coexistence between pinnipeds and marine industries must be achieved, but for this to occur wildlife populations must be protected while safeguarding marine livelihoods (*1, 18*). However, managing this triple bottom line requires a quantification of the economic, social and ecological impacts of pinniped-fishery conflict. Currently, we lack comprehensive analyses of the quantitative impact of pinnipeds on fisheries, preventing ecosystem-based management focussed on coexistence. More broadly, quantifying the unintended consequences of population recovery and conservation management will be critical as human populations and conservation initiatives expand.

Although there are widespread qualitative reports of pinniped depredation and damage to fisheries catch, the lack of a systematic quantitative global assessment of interactions between pinnipeds and fisheries remains a major barrier to better understanding fishery-pinniped conflict (*4, 15, 16*). Tixier *et al*. (*15*), for example, found that 214 distinct fisheries from 44 countries and all but two FAO fishing regions reported depredation from marine predators between 1979-2019, of which 30.8% were reports from pinnipeds. Marine predators were responsible for an 11% reduction in catch, although how much of this is attributable to pinnipeds is unknown These patterns are emphasised in targeted qualitative studies, where fishers overwhelmingly support management on pinnipeds (*19*). In Peru and Chile, one study reports 65% of surveyed fishers (n = 55) having interactions with pinnipeds (*20*), while another reports negative interactions with pinnipeds for the overwhelming majority (87%, n = 301) (*16*). For artisanal fishers in the Foça Monk Seal Pilot Conservation Area, Turkey, damage to fishing gear by the critically endangered Mediterranean monk seal (*Monachus monachus*) was reported 90 times (of 142 observations) between 1992-2004 (*21*). Increasing reports of pinniped-fishery conflict and the economic cost to fisheries highlights the need for more quantitative assessments of this global issue. Some reports of pinniped-fishery interactions pertain to biological interactions—concerns about impacts on recovering stock and food-web dynamics (*22, 23*). Conversely, there are reports of bycatch or fisher retaliation—describing the negative impacts of fisheries on pinnipeds (*24*–*27*). However, the metric that is most tractable to understand the economic impact of pinnipeds on fisheries is operational interactions, and specifically the frequency of interactions and damage to catch (*28, 29*). Operational interactions estimates are quantifiable and comparable, and thus well suited to synthesise this global issue.

To fully quantify the global impacts of pinniped-fishery conflict, we need to ascertain whether reports of pinniped-fishery conflict are representative. For instance, there may be reporting biases towards areas with a high degree of conflict, inflating estimates. To understand if this is the case, we need a predictive framework identifying areas where pinniped-fisheries conflict is expected to occur. A lack of interaction data in areas identified as potential hot-spots may indicate that other means of management such as legal/illegal culls have already been implemented by fisheries (*30*), thus highlighting areas where pinniped recovery has been slower or not occurred at all (e.g., for species that have become extinct such as *Zalophus japonicus*; (*31*)). Even if reporting bias is low, limitations of data collection and fishery management may mean interaction data is not available. Therefore, exploring spatial patterns in the reporting of operational interactions, and predicting where pinniped conflict may occur, is a vital step in quantifying this issue.

In the current study, we provide a quantitative global assessment of the impacts of operational interactions between pinnipeds and fisheries. First, we conduct a systematic literature review to synthesise comparable quantitative assessments of operational interactions of English literature from the Scopus and Web of Science databases. We use two negative operational interactions as dependent variables: 1) the proportion of normalised fishing days in which direct interactions between fisheries and pinnipeds were observed, and 2) the proportion of catch lost to pinnipeds. Interactions include any temporally explicit observation of pinniped activity in proximity to, or interference with, fisheries. We include only temporally explicit operational interactions, which were normalised against fishing effort in fishing days and collected primarily by independent observers, so that observations are comparable. Then, we use a mixed-effects meta-analysis to estimate pooled impacts of interactions and catch loss on fisheries and study heterogeneity. We explore how fishery size (large commercial vs. small artisanal/recreational), fishing gear type (net, line, trawl and mixed) and proximity to shore (near shore vs. offshore) influence operational interactions using meta-regression analyses.

Finally, we take key first steps in predicting the spatial extent of fishery-pinniped conflict, and investigate the spatial biases and global context of reported operational interaction data and their association with global fishing effort (*3*). The recent publication of global fishing data from Global Fishing Watch (*3*), provides an opportunity, when linked with pinniped occurrence, to begin addressing the global extent of fishery-pinniped conflict.

Our focus on quantitative operational interactions provides global evidence to inform economic and conservation policy, particularly those centred on compensation or conflict resolution schemes, whilst achieving ecological goals and protecting human livelihoods. We highlight the role of quantitative data in understanding interactions between fisheries and marine mammals and set out guidelines for future assessments of operational interactions.

## Results

Of 1,179 screened articles from Scopus and Web of Science, there were 376 publications focussed on coexistence and interactions between pinnipeds and fisheries, highlighting the global significance of this issue for fisheries. However, many articles presented qualitative assessments (n = 17), bycatch (n = 49), or investigated biological interactions (n = 48). We retained 36 articles that included quantitative and temporally explicit measures of operational interactions between pinnipeds and fisheries, both the proportion of fishing days with interactions (n = 23) and the proportion of catch lost (n = 22) (Figure 1). From the 36 retained studies, there were 33 observations for interactions and 33 observations for the proportion of lost catch. We normalised observations by the number of fishing days to account for differences in sampling effort and variance between studies. Retained studies reported observations of interactions over 2-20,693 fishing days (Figure 1b), with total observation periods of 25,867 and 6,929 days for interaction and catch loss data, respectively. An additional 11 studies presented surveys of fishers or managers giving qualitative measures of interactions and damage (Figure 1). These qualitative observations were excluded from the meta-analyses due to the omission of key variables required to standardise observations, including time period or use of qualitative versus quantitative estimates (e.g., qualitative categories used to describe frequency of interactions). However, in this survey data there are large proportions of fishers reporting regular interactions (mean = 64.8 %) and damage (57.8 %). The retained quantitative studies, while relatively sparse, are distributed across the globe, but there were general biases to South America and Northern Europe (Figure 1).

**Figure 1.**
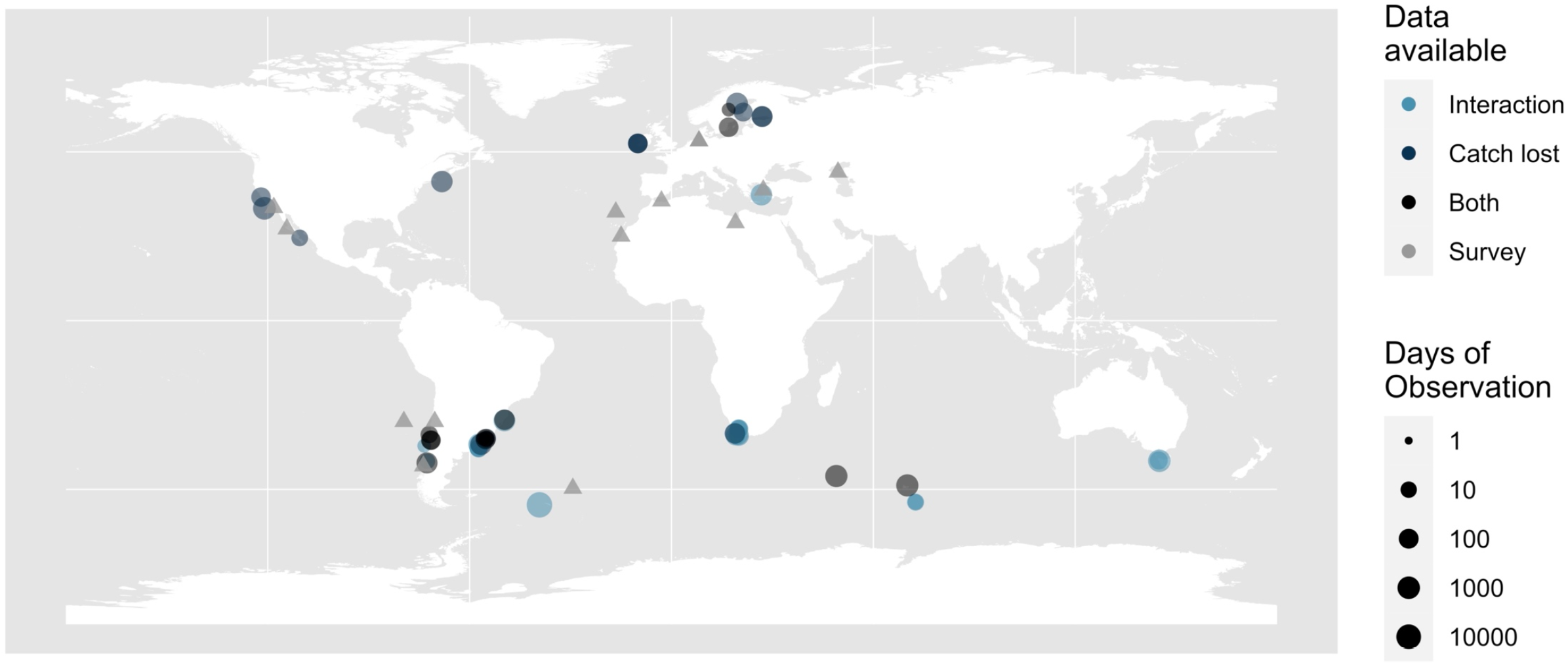
Global distribution of studies included in quantitative meta-analyses of operational interactions (n = 36), as well as qualitative survey studies (which were excluded from meta-analyses, n = 11) (b). The size of the points gives the number of fishing days in each study. The colour denotes the data available (type of operational interaction data) for each study: Light blue points indicate where interaction data were available, dark blue where catch loss data were available, black points where both interaction and catch loss data were available, and grey triangles indicate qualitative survey data that was excluded from meta-analyses.

We find evidence for extensive operational impacts of pinnipeds on fisheries, but a high degree of heterogeneity between studies (Figure 2). On average, 33.7 % [95 % confidence limits: 23.1; 46.3] of 25,867 fishing days were affected by direct interactions with pinnipeds (Figure 2a). Furthermore, 13.8 % [8.08; 22.5] of catch was lost to pinnipeds across 6,929 fishing days (Figure 2b). Together these results suggest that pinniped depredation and damage pose significant economic costs to fishery activity where interactions exist. However, the impacts of pinnipeds are not uniform across studies. This was accompanied by a high degree of heterogeneity in meta-analysis outcomes (Figure 2). For both negative interactions and damage, Cochran’s *Q* tests of similarity in binomial responses indicate highly significant differences in responses across studies (*Q* = 2140, p < 0.001 and *Q* = 997, p < 0.001, respectively), as well as substantial heterogeneity indices of *I*^2^, τ^2^, and *H* (Figure 2). This provides clear evidence that global operational impacts vary substantially and indicates that fishery or other characteristics may be crucial in driving differences in operational interactions.

**Figure 2.**
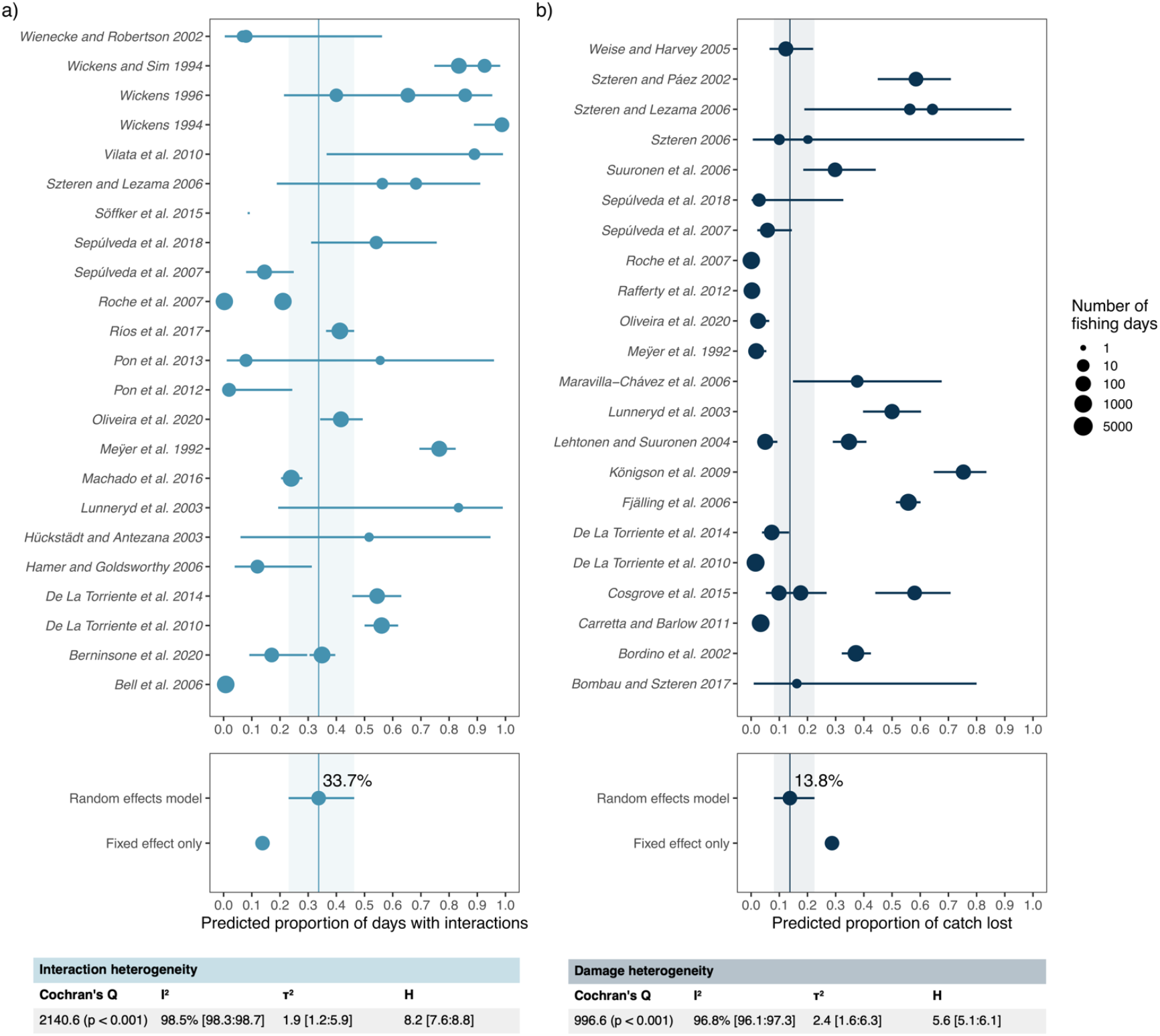
Operational interactions between pinnipeds and fisheries. Meta-analysis results (top panels), pooled estimates (middle panels) and heterogeneity measures (bottom panels) for studies quantifying the proportion of fishing days with interactions (a; n = 18) and the proportion of catch lost in those days (b; n = 19). In the top panels, each point is an observation from a retained study with estimated 95% confidence limits (Clopper-Pearson interval; Balduzzi et al. (*32*)), with the size of the point giving the number of fishing days. The middle panel gives the predicted pooled estimates for both the random effects model and fixed effects only model. The 95% confidence limits (shaded area) and pooled estimates (vertical line) are presented for the random effects model. The tables in the bottom panels are the heterogeneity measures, with 95% confidence limits.

Fishery characteristics predict the frequency and impact of operational interactions with pinnipeds (Figure 3). Fishery size (two-level categorical predictor variable) is reported in retained studies, and distinguishes between large-scale commercial fisheries, which typically operate at greater distances from shore, and small-scale/recreational fisheries typically operating closer to shore. Specifically, we find that small-scale/recreational fisheries using nets are significantly more likely to have interactions with pinnipeds. Note that as fishery size and location were highly correlated, we excluded location from meta-regressions. For both proportion of fishing days with interactions and proportion of catch lost, fishery size has a significant impact on operational interactions (*β* = 1.05 *±* 0.48 SEM, *z* = 2.17, p < 0.05 and β = 1.88 *±* 0.62 SEM, *z* = 3.02, p < 0.01, for interactions and damage, respectively; Figure 3a & Figure 3c). The predicted proportion of fishing days with pinniped interactions for large-scale fisheries is 21.2 % [11.3; 36.0], in comparison to 43.4 % [22.9; 66.4] for small-scale/recreational fisheries (i.e., small fisheries were over twice as likely to interact with pinnipeds). The proportion of damage was five times greater in small-scale/recreational fisheries, with a predicted damage of 21.8 % [7.62; 48.6] of catch compared to 4.86 % [1.50; 10.6] in large-scale fisheries (Figure 3a & Figure 3c).

**Figure 3.**
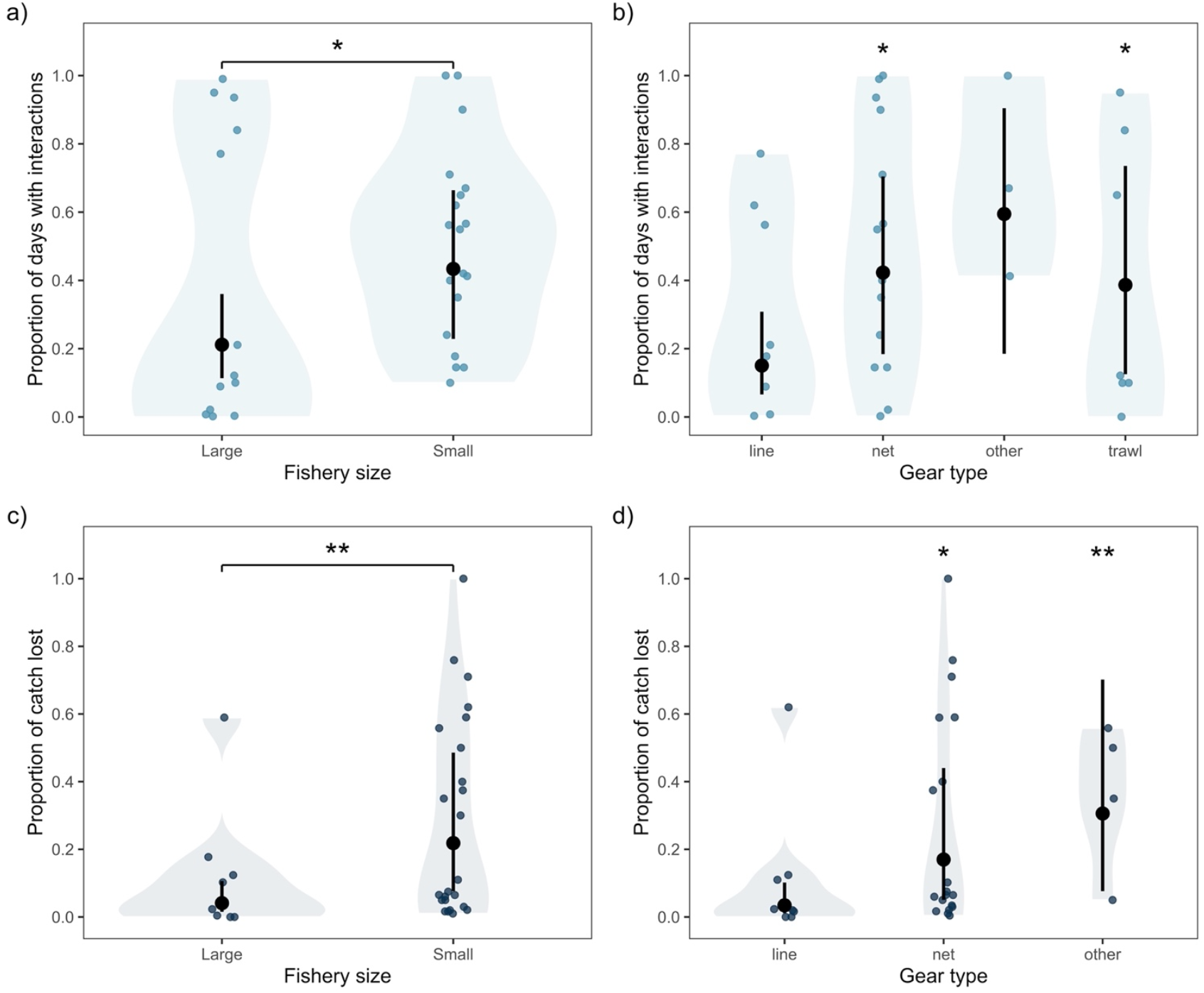
Fishery characteristics explain differences in operational interactions between fisheries and pinnipeds. The relationship between a fishery’s size (a & c) and the gear type used (b & d), and the proportion of fishing days with interactions (a & b) and the proportion of catch lost (c & d). Coloured points are observations from retained studies, with shaded areas indicating the approximate symmetrical data density. Black points and lines are the predicted mean operational interactions with 95% confidence limits. Significance displayed at the 95% level.

We find that the fishing gear used, namely net, line, trawl or other operations, predicts operational interactions. Fisheries using nets have a greater proportion of interactions and a greater proportion of catch lost (Figure 3b & Figure 3d). Net fisheries are predicted to interact with pinnipeds on 42.3 % [18.4; 70.4] of fishing days and lose 17.0 % [5.04; 44.0] of their catch, in contrast to 15 % [6.58; 30.8] of fishing days and 3.44 % [1.11; 10.2] of catch lost in line fisheries (Figure 3b & Figure 3d). Fisheries using various ‘other’ gear types and trawling also generally report a greater number of operational interactions (Figure 3b & Figure 3d), but reduced sample sizes in these groups prevents reliable inference. Other features of retained studies, including reporting source (e.g., academia, government) pinniped species, local pinniped population trend, and target fish species do not explain variation in operational interactions—although this could be due to low group-level or study-level sample sizes for these features.

To provide a first step in predicting spatial patterns of pinniped interactions and explore potential spatial biases in retained studies, we construct a composite spatial index at a resolution of 0.5°, of the potential for pinniped-fishery interactions from fishing effort (log10-transformed) (*3*), occurrence data for pinniped populations in IUCN category ‘Least Concern’, and the distance of each 0.5° pixel centroid from shore, with closer to shore areas indicating higher pinniped density due to haul out and breeding locations (Figure 4). The global index highlights nine regions that are expected to have the highest occurrence of pinniped-fishery interactions: the Bering Sea, the West-Coast of North America, the temperate and cape region of South America, the North-East of North America, Iceland, temperate and arctic regions of Northern Europe, the Cape of Southern Africa, Southern Australia and New Zealand, and the Seas of Japan and East China (Figure 4). The current study has quantitative observations from six of the nine regions, with data lacking from the Bering Sea, Iceland, and the Seas of Japan and East China. However, there is a current bias in study localities, particularly towards South America and Northern Europe, which comprised 23 of the 36 retained studies. Nevertheless, the average potential for pinniped-fishery interactions is substantially higher in locations where studies were carried out (Figure 4), suggesting that the measures of global fishing effort, non-threatened pinniped occurrence, and proximity to coast are useful proxies for predicting areas with operational interactions. Furthermore, these results suggest that retained studies capture much of global operational interactions.

**Figure 4.**
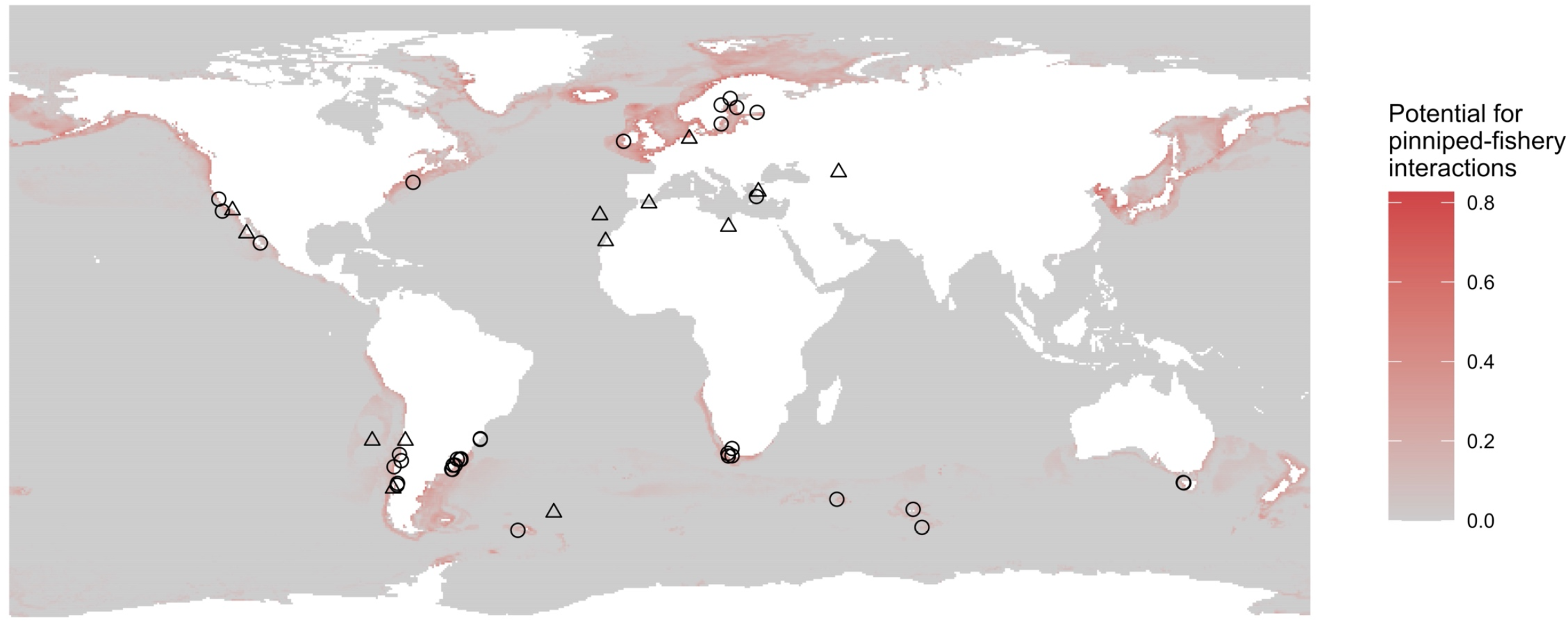
Composite map of the potential for pinniped-fishery interactions. Pixels give the composite index of the potential for pinniped-fishery interactions at a spatial resolution of 0.5°, for which high values (red) indicate high fishing effort, presence of non-threatened pinnipeds and proximity to coast. Black points give study localities, where open circles indicate quantitative studies (n = 36) retained in meta-analyses, and open triangles indicate qualitative studies (n = 11), which were excluded.

## Discussion

Fishing has occurred for at least 40,000 years, and conflict with marine predators such as pinnipeds is just as old (*33, 34*). Historical methods of resolving conflict between pinnipeds and fisheries ranged from hunting to systematic or managed culls, which led to drastically reduced pinniped numbers during the 18^th^ and 19^th^ centuries (*10*). Widespread introduction of marine mammal protective legislation in the 20^th^ century means previous conflict resolution strategies are largely unavailable to managers today. It is the very success of protective legislation that is exacerbating current conflict between recovering pinniped populations and fisheries. Today’s marine industries must operate alongside large populations of pinnipeds, which means fisheries managers and conservation scientists and practitioners must find new ways of managing conflict. These new conflict management strategies must balance the triple bottom line—ensuring a balanced outcome of social, ecological and economic objectives. However, there is a lack of understanding and consideration of this issue, preventing appropriate policies to overcome it (*4, 15*). We quantify the cost of conservation success by undertaking a global synthesis of the extent and severity of operational interactions between pinnipeds and fisheries. In areas where operational interactions are quantified and reported, fishers interact with pinnipeds on 33.7% of fishing days and report damages of 13.8% to total catch. These results highlight pinniped fishery conflict as a global economic issue that requires continued and extended management as pinniped populations continue to recover.

While operational interactions were globally widespread, the costs of conflict did not affect all fishers equally. We found that small-scale artisanal/recreational fishers were twice as likely to have interactions with pinnipeds on a given fishing day, and lose five times as much catch, compared to large-scale or commercial fisheries. These increases in operational interactions were also observed for fishing operations closer to shore. Additionally, fishers using nets were over three-times as likely to interact with pinnipeds, and lost over twice as much catch compared to line fisheries. Our results highlight that there are important equitable considerations in the management of pinniped-fishery conflict. Equitable conservation outcomes that protect the livelihoods of key stakeholders, are becoming increasingly important as human populations increase (*35, 36*). These results add to an increasingly dominant narrative in human-wildlife conflict research, that the poorest sectors of society often experience a disproportionate burden from conflict (*37*–*39*). For example, Fentaw *et al*. (*39*) found that pastoral communities in Southern Ethiopia experienced livestock losses of 37% due to wildlife in a neighbouring protected area. Thus, to ensure equitable outcomes, marine managers could tailor future management solutions to focus on conflict mitigation in areas with small-scale, net-based fishery operations.

In addition to a quantitative synthesis of the frequency and severity of operational interactions, we also take the first steps to predicting pinniped-fishery conflict worldwide. We find that operational interactions are more likely in areas where pinniped occurrence (*11, 40*) and fishing effort (*3*) overlap. Of the areas we identify as potential hotspots of operational conflicts, however we lack reports of interactions from the Bering Sea, Iceland, and the Seas of Japan and East China. Lack of studies from these regions could indicate a bias toward English publications, which were the only language of primary literature included in our study, or insufficient prey abundance to support pinniped populations due to overexploitation from fisheries (*41*). Finally, a lack of reporting in these areas may indicate that alternative conflict management strategies such as culls and hunting have already been implemented (e.g., (*31*)). One limitation to our spatial predictions is the use of global fishing watch data that is biased towards large-scale commercial fisheries with centralised GPS tracking. Given our finding that interactions are disproportionally experienced by small-scale fisheries, future predictions should incorporate spatial fisheries data from small-scale and artisanal operations (e.g., (*42*)). Better predictions of operational interactions will enable us to further illuminate the global extent of this conflict, highlight global reporting biases, and ultimately improve development of targeted management strategies.

There are a number of limitations that are necessary to acknowledge to improve future efforts to quantify pinniped-fishery conflict. First, despite widespread reporting (*15*), only a small proportion of identified studies report quantifiable estimates of damage or interactions to advise management. To inform conflict mitigation strategies, more temporally explicit quantifiable estimates of economic losses to fisheries are needed. Second, very few studies extend work on operational interactions to quantify the economic costs of pinniped conflict (but see (*43*)), most likely due to difficulty in estimating loss to catch that occurs under the surface. However, estimates of the costs incurred through loss/damage of fishing gear, which are only very rarely reported, may further advance understanding of this issue. Third, survey-based methods are prone to biases, highlighting the need for data collected by independent observers (*44*).

### Conclusion

Fisheries conflict with marine predators such as pinnipeds is a global issue, and management strategies must balance conservation, social, and economic outcomes. To balance this triple bottom line managers need information on the economic outcomes of conflict. We provide this critical missing element through a quantification of the impact on fisheries of operational interactions with pinnipeds. We find that pinniped interactions are common and damages significant. Equitable management will need to address the heterogeneous distribution of these costs, particularly as these costs are often disproportionately borne by smaller scale fisheries. Using this information, managers can begin to work towards solutions that balance ecological, social, and economic desires and outcomes for society. Coexistence with marine predators is the end goal, but knowledge of individual fishery-contexts will be key to achieving working solutions.

## Methods

Our approach was to identify relevant studies from the natural and social sciences using SCOPUS and Web of Science. We conducted an initial sorting of relevant studies based on title, abstract, and key words screening, published in all available years equating to 1^st^ January 1960 through 17^th^ August 2022 for SCOPUS, and 1^st^ January 1945 through 17^th^ August 2022 for Web of Science. The percent overlap of relevant papers between the two search platforms was calculated and the relevant papers were consolidated. We then developed a rubric to guide data extraction and completed a scoping study to determine whether sufficient data existed to calculate our effect sizes of interest. In what follows we provide an overview of our systematic literature review, the definition of the effect sizes of interest, the final rubric for data extraction, then our analytical methods.

### Literature review

We constructed a two-part search string. The first part focussed on fisheries human-wildlife conflict or variants thereof, including human-wildlife conflict, human wildlife interaction, HWC, operational interaction, conflict management, depredation, overexploitation, animal welfare, conservation conflict, marine mammal-fishery interaction, marine mammal predation, artisanal fishery, coastal fishery, commercial fishery, purse-seine fishery, gillnet fishery, interaction with fisheries, marine mammals and fisheries, seal-induced catch*, seal-induced damage*, poaching, culling, acoustic harassment devices, anti-predator nets. The second part used several terms covering groupings of pinnipeds to refine the breadth of search results. Specifically, we searched for pinnipeds, seal*, fur seal, true seal, sea lion*, walrus, marine mammal*, marine predators as key search terms. Based on our search string, we had 966 hits in SCOPUS and 188 unique hits in Web of Science. An initial assessment based on title and abstract considered that results were relevant if pinnipeds and fisheries were both mentioned. This left us with 376 papers that we reviewed in depth. We broadened this search through following up hits from reference list of relevant papers. This additional search method gave us 28 additional papers.

### Operational interactions of interest

We focus on two operational interactions in the current study, which were used as response variables in meta-analyses. We use operational interactions in which direct observations of pinnipeds interacting with/damaging fishery operations are made. We ignore indirect biological interactions for quantitative meta-analyses because biological interactions are both less comparable (relating to food-web differences) and harder to observe (underneath the surface) (*45*). Furthermore, we do not include observations of retaliation by fishers on pinnipeds, which are rarely quantified, or bycatch effects, which have been explored in detail (*25*–*27*). Therefore, our two effect sizes of interest centre around the likelihood that fishers will have interactions with pinnipeds, and the impact of these interactions on catch. Critically, in order to obtain comparable metrics for meta-analysis, we retain only temporally-explicit measures of operational interactions. The normalising temporal component was number of fishing days. The rationale for this measure as opposed to using fishing trips or sets was that these measures vary substantially between fisheries and gear type. While fishing days may also be biased where fishing activities are not constant throughout each fishing event, they provide a more comparable quantitative measure of the temporal component of fishing activities. For two studies (*28, 46*), we used (spatially) neighbouring studies with similar fishery size and practises to normalise fishing trips into fishing days. Furthermore, we primarily used studies in which independent observations were made on board fishing vessels, which reduces potential biases.

The first effect metric we calculate is the proportion of fishing days that have interactions with a pinniped species. An interaction in this context is broadly defined as the presence and activity of pinnipeds when fishing activities were taking place. These interactions can range from observations of pinnipeds swimming near (e.g. <50 m from vessels) fishing operations (*47*) to direct observations of pinnipeds damaging catch (*45*). We opted to group together these interaction modalities for the purpose of the study, as any less direct observations are likely an underestimate of interactions under the surface (*47*). Thus, the effect of interest was the proportion of reported fishing days in which interactions were observed.

The second effect metric that we calculate is the proportion of catch that is damaged by pinnipeds for a specific fishery and specific number of fishing days. This metric, where reported, is more explicit, for which observers have provided quantitative estimates of the proportion of catch lost to pinnipeds either through direct observations of damage, catch damaged (*e*.*g*. with bite marks; (*46*)), or through counterfactuals comparing sites with or without pinniped activity. In both cases, the sample size used in meta-analyses estimates is the number of fishing days.

### Covariates

To explore potential drivers of differences in operational interactions between studies, we collected a set of biological, fishery, and study characteristics. Biological characteristics included categorical variables of the focal pinniped species, the reported local population trend of the focal pinniped species, the target fish species, and a binary indicator of whether a study targeted multiple fish species. The local population trend of the focal pinniped species was obtained from information in the study itself and reported as decreasing, stable, or increasing. We did not use external sources such as IUCN regional assessments for this variable, because we aimed to capture the specific spatio-temporal context of pinniped population change. Fishery characteristics recorded included categorical variables of the fishery size (commercial [‘large’] or small-scale/artisanal [‘small’]), the location of fishing operations (near-shore or off-shore), the type of fishing gear used (net, line, trawl, or other), and binary indicators of reporting of pinniped prevention activities (*e*.*g*. use of audio exclusionary devices; (*48*)), economic compensation for losses due to pinnipeds, and retaliation of fishers towards pinnipeds. Fishery size was chosen to capture broad differences between commercial and non-commercial fisheries, for which fleet and boat sizes were combined with information in each study’s methodology. Other study characteristics included the study location and country, the source of the study (affiliation of the first author: academia, government, non-governmental organisation, fisheries, and unknown), and the method of data collection (survey, observations on board, cameras, and logbooks). The study location was retrieved from coordinate bounding areas (rectangular) reported in each study. Where maps and areas were not reported, a bounding area was estimated from reported coordinate locations, which were expanded to a bounding box by adding/subtracting 5 degree-seconds to each coordinate. The source of the study was collected from the affiliation of the first author, which indicated the employer of the person carrying out the majority of the work on research effort.

### Meta-analysis

We tested whether the proportion of fishing days with interactions and the proportion of catch lost were significantly different from 0 using binomial mixed-effects meta-analyses implemented in the *metafor* and *meta* R packages (*32, 49*). We used R version 4.1.3 for all analyses (*50*). Binomial meta-analyses were constructed using a successes and trials framework, for which trials were used to give the weighted sampling variance for each study. Thus, successes were calculated from the proportion of interactions and catch lost, given the total number of fishing days. Trials were the total number of fishing days. Models were fit using logit transformed observed proportions (*49*). We used both fixed-effects and random effects models to account for repeated observations from the same study (n = 4 and n = 2 studies for interaction and damage proportions, respectively). In meta-analyses, we calculated back-transformed confidence limits at the 95% percent level. To estimate the heterogeneity of operational interaction estimates, and thus how variable reported interactions were, we performed a set of heterogeneity assessments. These included the Cochrans *Q* test for heterogeneity, and the *I*^2^, τ^2^ and *H* indicators of heterogeneity. Generally, these methods give an estimate of the average distance of observed proportions to the model pooled estimates.

Following meta-analyses models, we performed meta-regressions to explore how biological, fishery, and study characteristics were associated with operational interactions. Meta-regressions were fit to the initial meta-analysis models using the *metareg* function of the *meta* package. To prevent overfitting due to low sample sizes from retained studies, we included only univariate meta-regression models where a single predictor of interest was included. Low sample sizes also prevented us from including pinniped species and fish species as predictors in meta-regressions. However, the variation in operational interactions between pinniped species and fish species was visually explored in the raw data.

### Spatial analyses

To explore the spatial biases in quantitative reporting of operational interactions and to predict where pinniped-fishery conflict may occur, we developed a composite spatial index of the potential for pinniped conflict. We constructed this index from three core components that are likely to be important for pinniped-fishery interactions, i) global fishing pressure, ii) the occurrence of pinniped populations in IUCN category ‘Least concern’, and iii) the proximity of each spatial unit of analysis to a shoreline. The overall rationale for this index was to capture areas where fishing activities overlap with pinniped occurrence, whilst accounting for the increased presence of pinnipeds closer to shore. This global index was constructed at a spatial resolution of 0.5° (approximately 55 km grid-squares at the equator) using the *rasterize* package (*51*). The index was weighted equally with mean global fishing effort from 2012-2020 (*3*), the spatial distribution of non-threatened pinniped species from the IUCN red list of threatened species (*11*), and the proximity of 0.5° cells to the closest shore (*40*) (Figure SX). For global fishing effort, the data from Global Fishing Watch, sourced from Kroodsma et al. (*3*), is biased towards commercial fishing operations and omits many small-scale/recreational activities. However, global spatial data on small-scale/recreational fishing effort is not currently available. Therefore, we use our global fishing effort as a proxy for both commercial and small-scale/recreational fishing activity. We calculated fishing effort in each grid square as the sum of fishing hours for all vessels operating in each grid square between 2012-2020, which was scaled to an index between 0-1, first standardising fishing hours (with one hour added to avoid observations of 0) on a log_10_ scale and dividing by the maximum sum of fishing hours across grid squares. Occurrence data from non-threatened pinniped species, listed as ‘Least-Concern’ on the IUCN red list of threatened species, was chosen to reflect species that are most likely to have negative impacts on fisheries due to higher population sizes (but see (*21*)). However, spatial overlap between threatened and non-threatened species was high. Occurrence was converted to an index of between 0-1, where occurrence was scored as 1 and non-occurrence as 0.01, to indicate a 1% probability of observing a pinniped in a given grid square.

## Notes

### Competing Interest Statement

The authors have declared no competing interest.

